# NanoCortex: A Unified Agentic System for Nanopore Sequencing Analysis

**DOI:** 10.64898/2026.05.19.726254

**Authors:** Qini Xia, Ziyuan Wang, Mina Shokoufandeh, Sara H. Rouhanifard, Meni Wanunu

**Affiliations:** Department of Bioengineering, Northeastern University, Boston, MA, USA; Department of Pharmacy Practice and Science, University of Arizona, Tucson, AZ, USA; Department of Physics, Northeastern University, Boston, MA, USA

## Abstract

Nanopore sequencing has enabled various layers of information about DNA and RNA sequence isoforms and chemical modifications. Yet, the archipelago of disjoint nanopore analysis tools makes navigating among these a significant challenge for the nanopore user. We present NanoCortex, a unified autonomous agentic framework designed to bridge this shortcoming by providing end-to-end data processing which ranges from raw signal basecalling to biological interpretation. Built upon Gemini API services that incur usage-based API costs and orchestrated through the Gemini Agent Development Kit (ADK), the system utilizes a multi-agent architecture to autonomously perform task parsing, code generation, iterative code-level self-correction of code, and scientific interpretation. Following code generation, the code can be used offline. Benchmarking reveals that NanoCortex achieves significantly higher usability across complex analytical tasks compared to general-purpose large language models. The framework seamlessly integrates experimental data with meta-analysis of publicly available, biological databases to facilitate the extraction of biologically meaningful insights from sequencing data without cumbersome computational steps.

## INTRODUCTION

Large Language Models (LLMs) have demonstrated significant utility across a diverse range of scientific disciplines^1-3^. This is exemplified by the emergence of domain-specific architectures, such as Med-PaLM^4^ and MedGPT^5^, which provides clinical support grounded in established guidelines, and BioGPT^6^, which excels in mining biological literature. Concurrently, the advent of DNA and RNA foundation models, such as DNABERT^7^, RNA-FM^8^, and Evo^9^, has revolutionized biological sequence understanding. By integrating diverse modalities, including genomic sequences and vast repositories of scientific literature, AI has evolved into an indispensable research tool. Moreover, the rise of agentic paradigms has further enhanced these capabilities by extending LLMs from simply answering questions to actively planning, querying, and executing tasks. This research helps connect current AI capabilities to the future goal of building more general, human-like intelligence that can advance science.

More recently, AI Agents have attracted significant attention in the field of bioinformatics as well^10^, including CellWhisperer^11^, which enables automated interpretation of multimodal single-cell data, and AutoBA^12^, which autonomously orchestrates bioinformatics workflows through tool-calling. Nanopore sequencing has emerged as a widely adopted long-read, native sequencing platform, valued for its ability to capture complex genomic and transcriptomic features in real time, however, it currently lacks a dedicated agentic AI ecosystem capable of mastering its unique analytical complexities. Oxford Nanopore Technologies (ONT) and the ONT community have developed a comprehensive repertoire of tools, ranging from basecalling (e.g., Nanopolish^13^, Guppy^14^, and Dorado^15^) and modification detection (e.g., Tombo^16^, Remora^17^, m6ABasecaller^18^, and Mod-p ID^19^) to signal processing and *de novo* transcriptome assembly (e.g., FLAIR^20^, StringTie2^21^, and Bambu^22^). Researchers can now access more complex genomic and epitranscriptomic architectures. Yet, this rapid diversification of these tools presents a significant challenge: selecting the optimal software and integrating datasets for specific biological questions remains difficult for biologists without informatics training. Beyond the standard demands of genomic analysis, nanopore sequencing presents distinct challenges due to the complexity of raw signal processing and the need to analyze long-read sequencing alongside epigenetic landscapes. Existing turnkey platforms like EPI2ME^23^ rely on domain knowledge in multiple displinaries including coding, molecular biology and genomics and lack the necessary flexibility due to the nature of nextflow pipeline. Meanwhile, conversational tools like ChatGPT^24, 25^ are limited to generating code and cannot functionally integrate with external tools to perform advanced and data-driven analysis.

Here we have developed NanoCortex, an autonomous agentic framework (Fig. 1), designed for the end-to-end Nanopore sequencing analysis workflows including basecalling, modification detection, *de novo* assembly, sequence analysis, DNA/RNA LLM-based function, and data-driven reasoning via integration of diverse biological knowledge resources. The code generated by NanoCortex is then downloadable and available for future implementation by the user. This approach provides a multi-scale integration of ONT sequencing information with biological phenotypes from publicly available sources.

**Figure 1.**
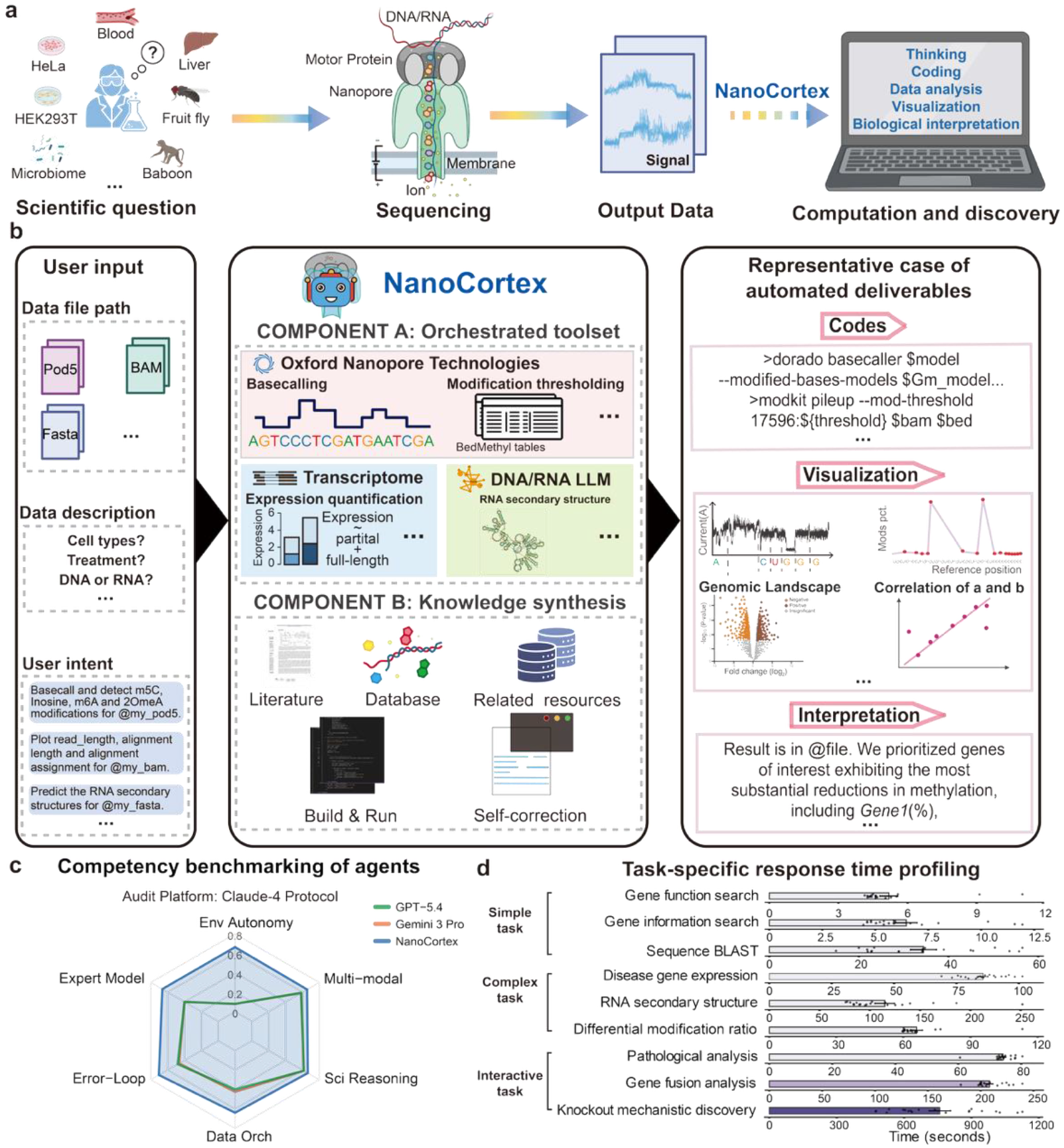
NanoCortex: A Unified Agentic System for Nanopore Sequencing. (a) Overview of the integrated nanopore sequencing and analysis workflow. The system uses computational analysis to bridge the gap between nanopore sequence data and analysis, visualization, and biological discovery/interpretation. (b) The end-to-end agentic workflow begins with a natural language query, in which the user specifies input file paths, data characteristics, and analytical objectives. The NanoCortex core comprises two parallel modules: Component A and Component B. Representative outputs are shown on the right, including generated code, visualization, and biological interpretation of the data. (c) Radar plot summarizing Claude-based evaluation across six capability domains: environment autonomy, expert model usage, error-loop correction, data orchestration, scientific reasoning, and multi-modal integration. Scores are derived from the aggregated functional completeness matrix. (d) Task-specific response time profiling. Response times across nine biological tasks (n = 20) utilizing Gemini 3 pro. Bars and error bars represent mean ± s.e.m., respectively. Individual points indicate raw replicates. Color intensity scales with mean RT. Some graphical elements in panels a and b were created using BioRender.com.

## RESULTS

### Nanocortex, an autonomous agentic framework for high-throughput ONT data

The standard long-read workflow for nanopore data encompasses sample extraction, library preparation, real-time sequencing, and data analysis (Fig.1a). Critically, the analytical phase requires proficiency in ONT-specialized software or custom scripts for data visualization and biological contextualization, creating a prohibitive bottleneck for both experimental biologists. To address this, we implemented an LLM-based agent capable of translating natural language queries into executable bioinformatic pipelines (e.g., the biologist tells the agent what it would like to test and the agent runs the test and reports the results).

NanoCortex is built upon a large language model–driven framework (Gemini)^26^ and orchestrated through the Gemini Agent Development Kit (ADK), enabling coordinated tool use and multi-step reasoning within a unified system (Fig. 1b; Supplementary Table S1; see Methods). As illustrated in Fig. 1b, NanoCortex operates through a structured workflow that transforms user input into automated analytical deliverables. Users provide a prompt together with data inputs (e.g., POD5, BAM, or FASTA files), as well as descriptions of the experimental design and analytical intent.

Inputs are processed within two coordinated functional modules (Orchestrated toolset and Knowledge synthesis). Orchestrated toolset integrates specialized tools for nanopore signal processing (Dorado^15^, Modkit^27^, Remora^17^), transcriptome analysis (StringTie^28^, FLAIR^20^), and RNA structure inference using RNAFM^8^, thereby establishing a unified environment for end-to-end sequencing analysis. Knowledge synthesis extends this capability by linking computational outputs to external scientific knowledge by incorporating literature resources, biological databases, and sequence similarity search (e.g., NCBI BLAST^29^). In addition, NanoCortex supports automated code generation and execution with built-in code-level self-correction, enabling automatic detection of execution errors, command revision based on error messages or tool documentation, and iterative workflow retry.

### NanoCortex evaluation compares with benchmark tools

While general-purpose models, including ChatGPT 5.4^24^ and Gemini 3.0 Pro^26^, implement subsets of nanopore analytical tasks such as basecalling parameter optimization or alignment troubleshooting, none of them provide a unified framework for end-to-end workflow execution. To systematically evaluate their capabilities, we performed an external competency audit, defined here as a structured, task-based evaluation across multiple operational dimensions. Each system was assessed with explicit scoring criteria, following LLM-assisted evaluation paradigms^30, 31^ (Fig. 1c; Supplementary Data 1).

The evaluation spanned key analytical tasks, including environment autonomy, expert model integration, agentic error correction, data orchestration, scientific reasoning, and multimodal output generation. Scores were assigned by Claude through a rubric-guided evaluation of software specifications and architecture-aware feature sets. Capabilities such as scientific reasoning were evaluated indirectly by human following the same standard. NanoCortex exhibited consistently high competency, forming a uniform performance profile under predefined rubric-based scoring criteria (See Methods and Supplementary Data 2). In contrast, online LLMs exhibited highly skewed capability profiles, with strengths in reasoning and multimodal interpretation but clear limitations in execution-centric dimensions such as environment-level autonomy, iterative error correction, and multi-tool orchestration (Fig. 1c).

To characterize the operational efficiency, we benchmarked the response time of NanoCortex across nine biological tasks, stratified into three tiers of increasing complexity: simple (basically one-subagent conducting task), complex (more than two subagents), and interactive (human-in-the-loop or agent self-revision). Quantitative assessment revealed a strong correlation (R = 0.952, Pearson) between task complexity and computational latency, with mean response times (RT) scaling from approximately 5s in simple tasks to over 800s in interactive workflows (Fig. 1d; Supplementary Fig. 1; Supplementary Data 2). While simple and complex tasks exhibited low RT, interactive tasks demonstrated higher performance variance due to API request exhaustion.

### LLM-based streamlined workflows connecting heterogeneous ONT software

To evaluate NanoCortex we deployed the agent across a series of complex analytical workflow, ranging from raw signal processing to deep-learning-based structural modeling (Fig. 2; Supplementary Data 3–9). The agent functions as a unified computational interface, successfully bridging various software that typically require disparate expertise and manual data formatting.

**Figure 2.**
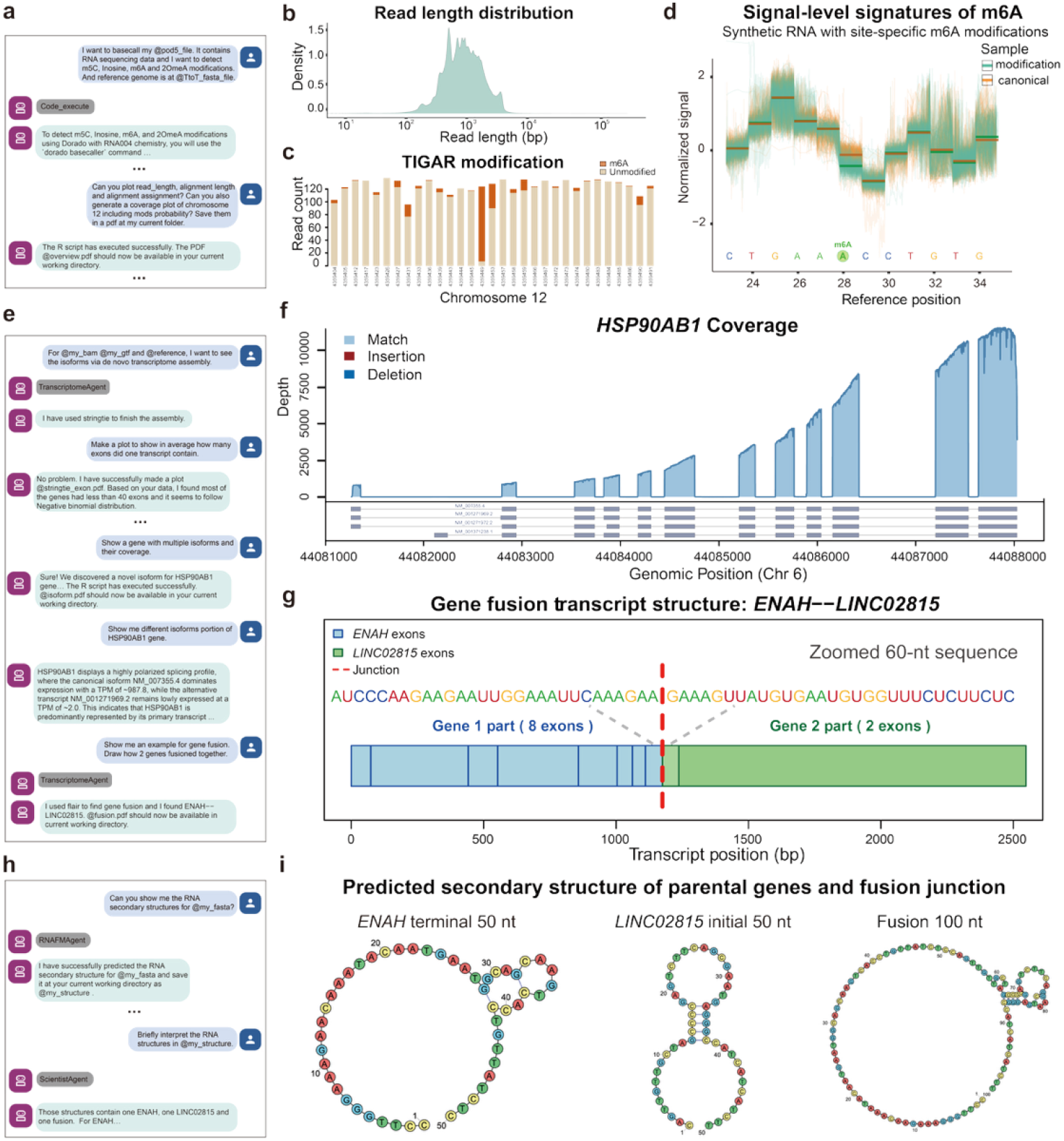
Agent-mediated autonomous analysis across diverse biological contexts. (a) Textual user input (blue text bubbles) for executing autonomous workflows, including basecalling and multi-modal modification detection for m5C, Inosine, m6A, and 2-Ome-A, as well as textual Nanocortex feedback (green text bubbles). Sub-agent selection by Nanocortex is shown in grey text bubbles. (b-d) Example output figures from Nanocortex-generated “overview.pdf” output, which include empirical density distribution of sequencing read lengths (b), quantitative mapping of RNA modification occupancy across specific genomic coordinates on Chromosome 12 (c), and normalized raw signal signatures distinguishing m6A-modified bases from canonical counterparts, where individual traces are shown in green (modified) and orange (canonical), with solid lines indicating the mean current at each position (d). (e) Textual input request to resolve isoform diversity and identify complex gene fusion events. (f-g) Example output figures from “fusion.pdf”, showing transcriptomic coverage and isoform diversity for the *HSP90AB1* gene, as well as the distribution of match, insertion, and deletion events (f). Structural architecture of the *ENAH*—*LINC02815* spliced fusion transcript, highlighting the junction between the two fusion partners (g). (h) Textual input request to predict and interpret RNA secondary structures from sequence data. (i) Predicted secondary structures of representative RNA regions, including the *ENAH* terminal 50 bp, the *LINC02815* initial 50 bp, and the 100 bp fusion transcript derived from these two segments.

We first tested the agent’s ability to conduct the primary ONT analysis pipeline. Upon receiving natural language instructions, the agent autonomously executed Dorado^15^ basecalling, Minimap2 alignment^32^, modification detection (m5C, pseudouridine, inosine, m6A and 2-Ome-A), read length distribution, site-specific modification landscapes, and subsequently generating plots for signal comparison (canonical vs. modification) (Fig. 2a–d; Supplementary Fig. 2). This pipeline, which proceeded from raw pod5 files as inputs to epitranscriptomic profiles as outputs, was achieved without human intervention, with modification landscapes in Fig. 2c restricted to sites supported by at least 20 reads (a filter we imposed).

We further extended the agent’s scope to complex transcriptomic assembly and fusion gene detection based on conversation (Fig. 2e). By coordinating Stringtie^28^, FLAIR^20^, and custom R-based visualization scripts, the agent identified isoforms across multiple gene loci, including *HSP90AB1* and additional representative genes (Fig. 2f; Supplementary Fig. 3). As summarized in Fig. 2f, StringTie resolved four transcript isoforms with high-confidence read support, NM_007355.4 (1,684 reads), NM_001271969.2 (1,097 reads), NM_001271972.2 (1,069 reads), and NM_001371238.1 (56 reads), with read counts providing quantitative support for isoform-specific expression. In addition, our framework identified an *ENAH*–*LINC02815* spliced fusion candidate, supported by 21 fusion-spanning reads, alongside 94 reads mapping to *ENAH* and 33 reads mapping to *LINC02815*. (Fig. 2g). The agent not only executed the assembly and gene fusion detection software via Stringtie^28^ and FLAIR^20^ respectively but also provided automated statistical interpretations of isoform dominance and alternative splicing events.

Finally, we integrated RNA-FM^8^ and RiboSketch^33^ to resolve the secondary structures of transcripts associated with the identified fusion event. Specifically, we modeled the terminal 50 nt of the upstream gene, the initial 50 nt of the downstream gene, and the 100 nt sequence spanning the fusion junction, as these sequence lengths were sufficient to resolve the local structural features surrounding the breakpoint. Structural predictions revealed a marked reorganization at the junction, with the loss of a loop structure observed in the parental transcripts (Fig. 2h–i). Consistent patterns were observed at extended sequence lengths (400 nt; Supplementary Fig. 4), supporting the robustness of the predicted structural reorganization. Notably, this structural reorganization prediction was not anticipated a priori and was revealed through the framework’s integrative analysis of sequence and structural features.

### NanoCortex establishes an iterative alignment framework for uncovering unreferenced biological sequences

Conventional reference-based alignment approaches map sequencing reads to a predefined genome reference through sequence similarity, but lack the ability to trace the biological origin of unaligned reads. NanoCortex introduces a novel paradigm for genome alignment analysis, utilizing an automatic-updating reference genome and an iterative alignment pipeline (Fig. 3a). Basecalled reads are first aligned to an initial reference genome, separating aligned and unaligned reads for downstream analysis. Unaligned reads are iteratively subsampled and searched against external databases to identify novel sequences, which are incorporated as supplemental references to update the genome and progressively recover previously unmapped reads. By addessing the constraints of fixed genomic templates and integrating real-time BLAST-mediated updates, this approach effectively uncovers biologically significant information often neglected by standard analytical pipelines, providing a higher-resolution assessment of transcriptomic integrity. As a proof of concept, we applied this automated workflow to the HEK293T cell line (Fig. 3b). Allowing updates to the reference genome uncovered 8,454 reads from Adenovirus 5 (Ad5) and 3,993 reads from SV40 Large T-Antigen (SV40) 3993 reads, which accounted for 0.084%, and 0.040% respectively in total reads (Supplementary Fig. 5). Reflecting the constitutive integration of Ad5 E1 and SV40 Large T-antigen in the HEK293T lineage, the successful identification of these viral signatures by our pipeline serves as a robust internal benchmark, confirming the superior sensitivity and feasibility of our pipeline^34-37^. This approach enables rapid identification of unknown contaminants or novel pathogens, while revealing unrecognized sources of contamination (e.g., mycoplasma), thereby enhancing experimental robustness and reproducibility.

**Figure 3.**
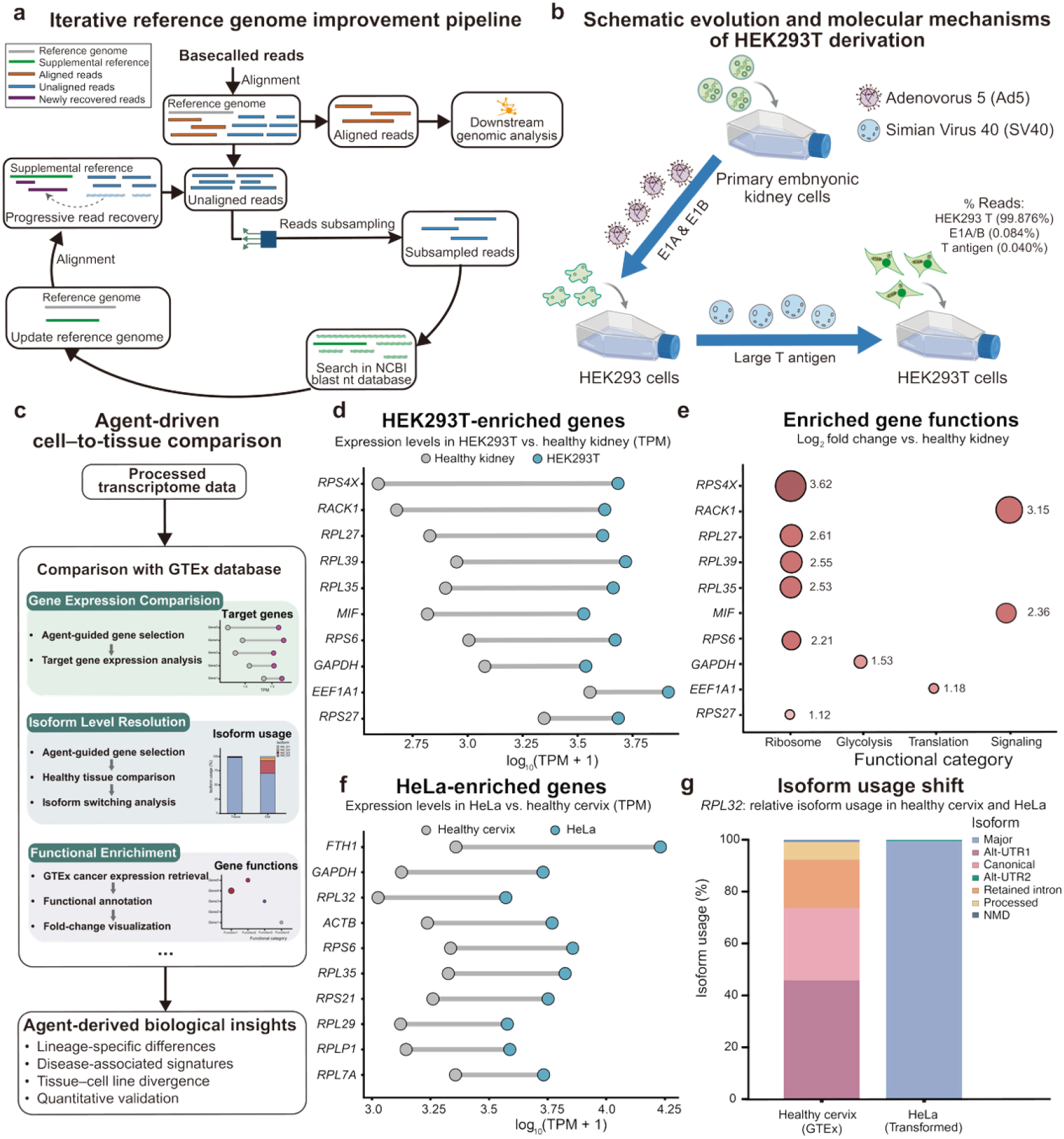
Agentic comparative transcriptomics and reference refinement across transformed cell lines. (a) The Iterative Reference Genome Improvement Pipeline identifies unaligned reads and performs BLAST searches to dynamically update the reference genome. (b) A schematic illustrates the details of the HEK293T transformation of primary embryonic kidney cells via Adenovirus 5 (Ad5) and Simian Virus 40 (SV40). (c) The agent-driven cell-to-tissue comparison pipeline integrates GTEx database to perform isoform-level resolution, gene expression comparison, and functional enrichment. (d) Expression levels of representative genes randomly selected from HEK293T-upregulated genes (n = 10), compared with healthy kidney tissue (GTEx). Values are shown as log_10_(TPM + 1). (e) Functional enrichment of HEK293T-enriched genes. Bubble size and numerical labels indicate log_2_ fold change relative to healthy kidney tissue, with categories including ribosome, glycolysis, translation, and signaling. (f) Expression levels of an additional set of representative genes, shown as in (d), confirming consistent upregulation patterns in HEK293T cells. (g) Isoform usage shift in *RPL32*, showing relative isoform proportions in healthy cervix (GTEx) and HeLa cells. Some graphical elements in panels b and c were created using BioRender.com.

### NanoCortex reveals reference-based comparative transcriptomic patterns across biological conditions

To expand the biological and clinical knowledge scope of NanoCortex, we integrated bulk RNA-seq data from the Genotype-Tissue Expression (GTEx)^34^ project to provide a reference atlas of gene expression across human tissues (Fig. 3c). By querying candidate genes, the platform quantified gene expression and identified differential transcript usage. Beyond individual gene-level comparisons, NanoCortex facilitates the automated Gene Oncology functional enrichment analysis of differentially expressed genes, enabling the identification of specific biological processes^38^. To evaluate the ability of the agent-driven framework to perform cross-context biological interpretation, we analyzed two wild-type cell lines, HEK293T cells and HeLa cells, in comparison with healthy GTEx tissues. Comparative analysis of HEK293T cells against healthy kidney tissue revealed a consistent upregulation of a subset of highly expressed genes (Fig. 3d). These genes, including ribosomal proteins (e.g., *RPS4X, RPL27, RPL39*) and translation-associated factors, exhibited elevated expression levels in HEK293T, indicating a shift toward enhanced biosynthetic activity. Functional annotation of these upregulated genes using the agent-driven pipeline further demonstrated enrichment in pathways related to ribosome function, translation, and signaling (Fig. 3e). A similar analysis in HeLa cells compared to healthy cervix tissue identified a distinct set of upregulated genes (Fig. 3f). To further investigate transcript-level regulation, we examined isoform usage across representative genes. Notably, genes such as *RPL32, RPL29*, and *GAPDH* exhibited a pronounced shift toward dominance of a single isoform in transformed cells relative to healthy tissue (Fig. 3g; Supplementary Fig. 6). ^39, 40^ These data demonstrate that the agent-driven framework enables systematic integration of transcriptomic profiling with physiological reference data, supporting scalable, cross-condition analysis.

### NanoCortex reveals structure-associated A-to-I editing patterns in transcripts

To explore the relationship between RNA modification and secondary structure, we first characterized the distribution of A-to-I editing across target transcripts (Supplementary Fig. 7). In *NDUFC2*, a pronounced enrichment of inosine was observed within the 3′ UTR, with editing levels ranging from 3% to 22%, whereas little-to-no editing was detected in other regions (Fig. 4a), consistent with previous reports^41^. Sequence analysis revealed that this region contains two complementary *Alu* elements, suggesting the formation of a double-stranded RNA (dsRNA) structure. Consistent with this, structural prediction using RNA-FM identified a distinct central duplex region within *NDUFC2* (Fig. 4b, blue box). This double-stranded architecture serves as the requisite substrate for ADAR-mediated A-to-I editing, encompassing a specifically targeted adenosine residue. Our structural characterization of the *NDUFC2* duplex is consistent with prior work defining the RNA structural requirements for ADAR occupancy^35, 36^.

**Figure 4.**
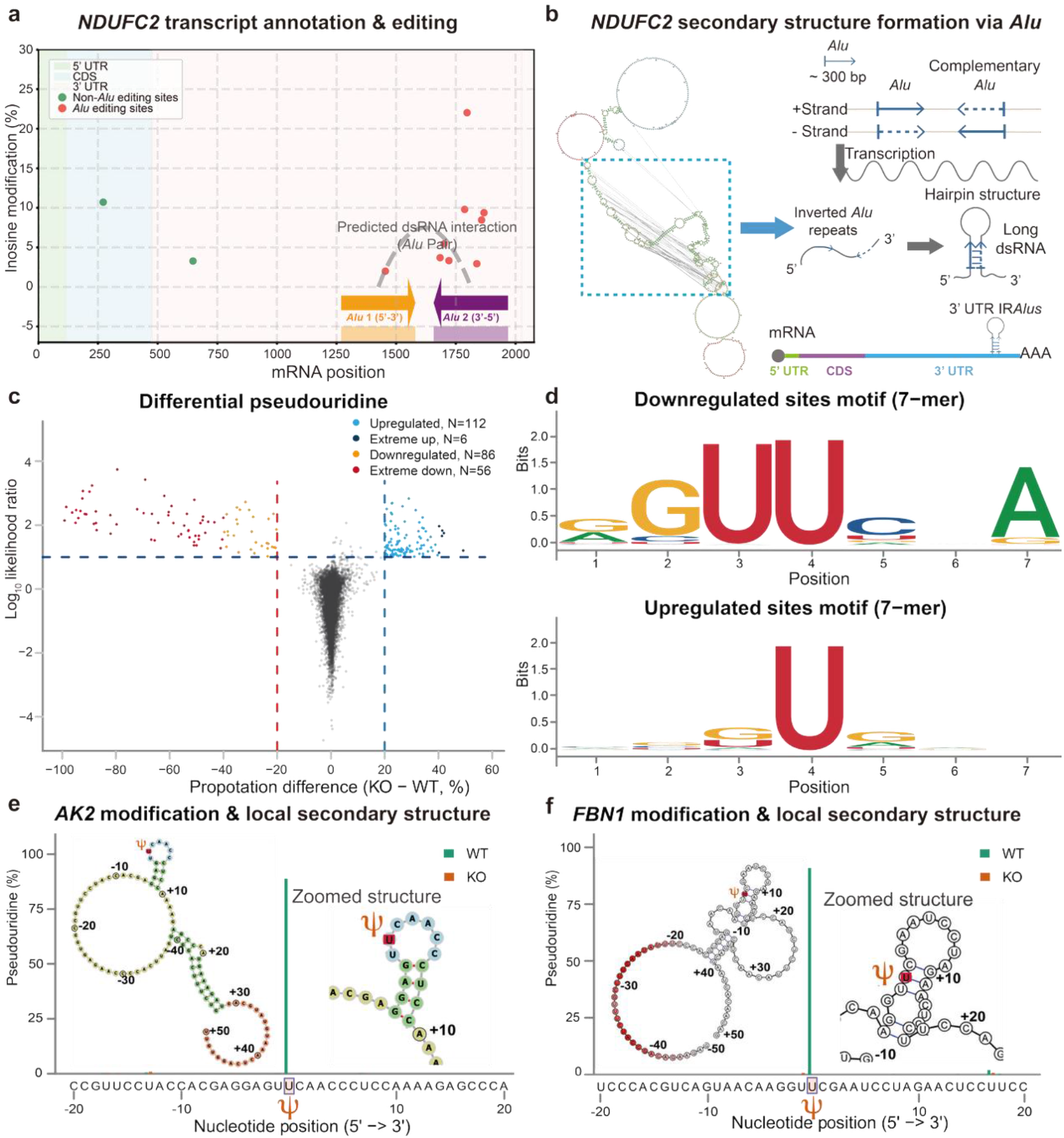
High-resolution mapping and structural modeling of RNA modifications. (a) Mapping of Inosine modification percentages across the *NDUFC2* transcript in HEK293T cell line. (b) Predicted secondary structure of *NDUFC2* and a schematic demonstrating how inverted *Alu* repeats facilitate the formation of a long double-stranded RNA (dsRNA) hairpin structure, serving as a substrate for inosine editing. (c) Volcano plot characterizing differential pseudouridine (Ψ) modification levels between wild-type (WT) and knockout (KO) conditions. Gray points represent a randomly sampled 0.1% subset of sites for visualization. (d) Sequence logos showing primary motifs of downregulated and upregulated Ψ sites (7-mer context). Letter height represents information content (bits) at each position. (e-f) Site-specific pseudouridine (Ψ) occupancy profiles and local secondary structures for the *AK2* and *FBN1* transcripts. The central position denotes the queried site, with flanking nucleotide positions shown (5′→3′). Insets display the corresponding local secondary structures.

### NanoCortex reveals sequence- and structure-dependent pseudouridylation patterns associated with *TRUB1*

We further evaluated the platform’s capacity to characterize epitranscriptomic profiles by comparing wild-type (WT) and *TRUB1* knockout (KO) HeLa lineages, where TRUB1 is a pseudouridine synthase responsible for RNA uridylation^37^. Differential modification ratio analysis revealed a global redistribution of pseudouridine (psi), with downregulated sites (i.e., lower overall pseudouridine proportions detected) showing greater effect sizes than upregulated sites (i.e., higher overall pseudouridine proportions detected), indicating that loss of *TRUB1* is associated with lower pseuroduridine modification levels (Fig. 4c). Analysis of primary sequence constraints via motif logos (Fig. 4d) revealed that downregulated pseudouridylation sites are significantly enriched for a degenerate GU[U]CNA consensus, in agreement with previously reported *TRUB1* target motifs^37, 42^. In contrast, upregulated sites exhibited no discernible motif preference, suggesting an alternative^37^.

We also recapitulated that *TRUB1* depletion suppresses psi modifications within stem-loop architectures via predicting RNA secondary sructures^37^. To validate the biological relevance of *TRUB1*-mediated pseudouridylation, we performed site-specific analyses of transcripts exhibiting strongly reduced pseudouridylation upon *TRUB1* knockout. Among these candidates, we highlight Adenylate Kinase 2 (*AK2*) as a representative example^43, 44^ which has been shown to exist in primary and immortalized human cells. NanoCortex identified a high-confidence pseudouridine site within the AK2 coding region that exhibited a marked reduction in modification occupancy following *TRUB1* deletion (Fig. 4e). Structural modeling indicates that this target uridine is positioned within the apical loop of a stem–loop structure (Fig. 4e; Supplementary Fig. 8), consistent with previous reports of TRUB1-associated stem–loop recognition motifs.^45^ Extending beyond this representative case, similar site-specific patterns were observed across additional transcripts, including *FBN1* and other examples (Fig. 4f; Supplementary Fig. 8), where pseudouridine sites exhibiting reduced occupancy in the knockout are located within similarly structured RNA regions.

## DISCUSSION

The development of NanoCortex represents a paradigm shift from conversational AI to autonomous agentic systems capable of performing complex nanopore sequencing analysis and code generation that is available for the user in future analysis needs. Through an execution-based, externally evaluated framework, we show that hierarchical multi-agent architectures are important for enabling robust end-to-end workflow execution. Specifically, NanoCortex autonomously diagnose faults via error logs and audit parameter requirements through help flags, establishing a robust self-correction framework that ensures reliable workflow execution.

Our framework integrates both computational pipelines and external knowledge resources, including database querying and literature retrieval beyond providing a collection of sequencing analysis tools. Importantly, we evaluated the system’s capacity for autonomous self-correction. The framework demonstrates robust error recovery without human intervention, including the identification and resolution of missing dependencies, incorrect file paths, and improper parameter configurations. This capability is facilitated by an internal orchestration harness that coordinates tool execution and iterative refinement, ensuring reliability in complex, multi-step analytical workflows. In terms of response time, we observed that task complexity is a primary determinant of execution latency. More complex tasks require extended reasoning and iterative querying, as well as increased computational overhead for downstream data processing, collectively contributing to longer response times compared to simpler tasks.

Beyond computational execution, we found the necessity of context-aware AI in elucidating biological signatures typically obscured by traditional bioinformatics pipelines. The discovery of hidden viral integrations (Ad5 and SV40), together with the identification of transformed-cell-associated gene expression divergence and systematic isoform usage shifts relative to healthy GTEx baselines^34^, demonstrates that the agent can generate biologically meaningful insights beyond routine data processing in an accessible manner.

Crucially, NanoCortex provides reasoning into the relationship between sequence, structure, and chemical modifications in RNA. The framework facilitates the per-site integration of epitranscriptomic modifications with their proximal RNA structural determinants. By systematically associating ADAR-mediated A-to-I editing with double-stranded regions and TRUB1-mediated pseudouridylation with specific stem–loop architectures, the system demonstrates a robust capacity to unify sequence and structural features for the comprehensive interpretation of epitranscriptomic landscapes.

As autonomous agents integrate into the laboratory, a better harness is required to further optimize agentic AI frameworks for high-throughput Nanopore sequencing workflows. Currently, the practical utility of such systems is constrained by LLM API access, response times, genomic data privacy, and token limits. Furthermore, this framework should extend from bioinformatics analysis to the direct management of MinKNOW control software enabling agents to autonomously modulate hardware parameters in response to live data streams during real-time sequencing.

## METHODS

### Overview of the NanoCortex framework

Nanocortex integrates large language model (LLM) reasoning with structured tool interfaces to dynamically construct and execute customized analysis of pipelines. The agent decomposes high-level analytical requests into structured tasks, retrieves relevant analytical modules, and assembles them into executable workflows. By translating natural language objectives into autonomous computational processes, the framework enables end-to-end analysis encompassing basecalling, modification detection, de novo assembly, sequence analysis, RNA representation learning, and biological interpretation through integration of external knowledge resources and databases for DNA and RNA molecules. A reproducible installation workflow is provided in Supplementary Table 2, with detailed command-line instructions available in the GitHub repository.

NanoCortex is orchestrated through the Gemini Agent Development Kit (ADK), with detailed implementation and task orchestration documentation available at https://google.github.io/adk-docs/tools-custom/performance/. NanoCortex relies on commercial Gemini API services and therefore require external service access and associated usage costs. Additional information regarding Gemini API access and pricing is available through the official Google AI documentation (https://ai.google.dev/gemini-api/docs/pricing).

NanoCortex adopts a modular architecture designed to streamline nanopore data analysis through task interpretation, orchestration, and execution. The system consists of four primary modules: (1) **Task Parsing Module**, which translates natural language queries into structured analytical objectives and selects appropriate specialized sub-agents; (2) **Nanopore Analysis Core**, which manages domain-specific processing, including basecalling, modification detection, transcript analysis (e.g., de novo assembly and gene fusion analysis), RNA representation learning, and related sequence-level and structure-level analyses; (3) **Code Generation Module**, which converts high-level workflow plans into robust executable command-line scripts; and (4) **Scientific Interpretation Module**, which synthesizes analytical outputs into biologically meaningful insights using LLM-guided reasoning.

### Task parsing and intent recognition

To facilitate autonomous task parsing and intent recognition, we developed a hierarchical multi-agent architecture composed of a root planning agent and specialized sub-agents. We used a root agent as a planner employing high-level prompts to evaluate user’s tasks and decompose them into discrete, manageable sub-goals. These sub-goals are subsequently dispatched to sub-agents. Like the root agent, each subagent used a similar pattern to understand the instructions generated by the root agent to select the necessary tools and execute the task (Supplementary Note).

### Orchestrated toolset

#### Workflow of Orchestrated toolset

The Orchestrated toolset manages domain-specific analytical workflows for nanopore sequencing data. Based on structured task definitions generated during query parsing, the system constructs analysis of pipelines composed of sequential processing stages. These workflows typically include signal processing, read alignment, feature detection, and downstream summarization.

Task dependencies between analytical stages are automatically resolved to ensure that workflows are assembled in a logical consistent order. This orchestration mechanism enables the system to dynamically generate analysis of pipelines tailored to the requirements of each user’s query.

#### Orchestrated toolset modules

The analytical framework integrates multiple processing modules that support an expanded range of nanopore sequencing analyses enabled by LLM-guided workflow construction. These modules include signal-to-sequence conversion, read alignment, DNA/RNA modification detection, transcript reconstruction, de novo assembly, gene fusion analysis, RNA representation learning, RNA secondary structure prediction (visualized by RiboSketch^33^), and splice-site prediction for transcript isoforms.

Each module is defined through structured interfaces describing its expected inputs, outputs, and execution requirements. This abstraction allows the agent to dynamically select and combine compatible analytical modules when constructing workflows. By representing analytical tools as standardized modules, the system maintains flexibility while ensuring compatibility between successive stages of the analysis pipeline.

In the current implementation, these analytical stages are performed using widely adopted nanopore analysis tools, including **Dorado**^15^ for basecalling, **minimap2**^32^ for read alignment, **Modkit**^27^ for RNA modification detection, **FLAIR**^20^ for transcript isoform and fusion analysis, **Stringtie**^28^ for transcript assembly and quantification, **Remora**^17^ for nanopore signal-level analysis and visualization, and **RNA-FM**^**8**^, a large-scale RNA foundation model, for generating sequence embeddings that support downstream structural and functional inference. A comprehensive list of computational tools is provided in Supplementary Table 1^46^.

### Knowledge synthesis

#### Workflow of knowledge synthesis

The Knowledge Integration module enables biological contextualization of computational outputs by connecting analytical results with external biological databases and knowledge resources. This module manages database connectivity and coordinates retrieval of relevant biological information required for downstream interpretation.

Following task parsing, the system identifies the appropriate specialized sub-agent responsible for the requested biological analysis. Based on the structured task representation, the agent retrieves compatible analytical tools and constructs interpretation of workflows tailored to the query objective.

These workflows may include sequence similarity searches, transcriptomic dataset comparison, RNA modification annotation retrieval, and external knowledge discovery through web-based resources.

#### Processing modules

To support these analyses, NanoCortex integrates multiple biological databases and computational resources. These include NCBI BLAST (https://www.ncbi.nlm.nih.gov/blast/)^29^ for sequence similarity searches, Google Search (https://www.google.com/) for general knowledge discovery, PubMed (https://pubmed.ncbi.nlm.nih.gov/) for literature retrieval, GTEx (https://gtexportal.org/)^34^ for transcriptomic expression data queries, and MODOMICS (https://iimcb.genesilico.pl/modomics/)^47^ for RNA modification annotation.

NCBI BLAST^29^ identifies homologous sequences through similarity searches against reference databases. Google Search retrieves publicly available biological knowledge and web-accessible resources related to genes, RNA molecules, or functional annotations. PubMed provides access to relevant scientific literature associated with specific genes, RNA structures, or modification types. GTEx supplies transcriptomic expression profiles across human tissues, enabling contextual interpretation of gene-level observations. MODOMICS provides curated annotations of RNA modification types and associated biochemical pathways.

#### Code generation, execution, and self-correction

We implemented an autonomous code generation module facilitated by the Gemini ADK to translate high-level intent into executable computational logic. Upon receiving a partitioned sub-task, the agent performs a semantic mapping of natural language prompts against a curated database of software-specific command protocols. This process integrates these domain-specific requirements to produce task-optimized scripts across Python, R, and Bash environments. By grounding the generative process into a specialized command database, the architecture ensures that the generated code maintains strict adherence to the syntax and parameter constraints of the underlying bioinformatic tools.

We developed a robust execution tool that processes sequential commands within a controlled shell environment. This module facilitated the serial execution of Bash, Python, or R commands. Prior to execution, all synthesized code undergoes a mandatory security audit by a Reviewer Agent. This secondary supervisory layer performs a static analysis of the proposed commands to identify and filter potentially unsafe operations or unauthorized system calls, thereby maintaining a secure and deterministic computational state. Moreover, the system monitors return codes in real-time, and any non-zero exit status triggers an immediate halt to prevent downstream error propagation. If the system had the error information, the agent leverages captured error logs to autonomously diagnose the underlying fault. To refine its understanding of tool-specific syntax, the agent autonomously invoked help flags (e.g., --help) to audit parameter requirements and iteratively regenerate the command sequence. This closed-loop refinement continues until the system achieves functional convergence, ensuring that the final output aligns with the user’s primary research objective.

### Benchmarking and performance evaluation

To systematically evaluate the analytical capabilities of NanoCortex, we designed a set of benchmark tasks representing common nanopore RNA analysis scenarios, including basecalling, RNA modification detection, transcriptome assembly, isoform quantification, and biological interpretation.

NanoCortex was benchmarked against state-of-the-art AI systems, including Gemini 3 Pro and ChatGPT 5.4. Rather than relying on predefined binary scoring, we implemented an external competency audit framework to evaluate system-level capabilities. Specifically, each system was assessed by an independent auditing agent (Claude, https://www.anthropic.com) using a standardized prompt that defined evaluation criteria across key dimensions, including environment autonomy (e.g., file system access, environment setup, and GPU utilization), expert model integration (e.g., RNA foundation model usage), agentic error correction (e.g., iterative loop-back and traceback-driven repair), data orchestration (e.g., multi-format handling and database integration), scientific reasoning (e.g., motif discovery and differential analysis), and multimodal output generation (e.g., visualization and report synthesis). The full task list, scoring criteria, and audit outputs are provided in Supplementary Data 2.

Scores were assigned by the auditing agent based on documented system capabilities, architectural constraints, and the degree of autonomous support expected for each benchmark task. The scoring ranged from 0.0 (architecturally infeasible) to 1.0 (fully supported), with intermediate values reflecting partial capability, execution gaps, or dependence on external human intervention. This framework was designed to assess system-level operational competency beyond static module-level evaluation.

To further characterize operational performance, we evaluated the response time (RT) of NanoCortex under different underlying large language model backends, including Gemini 3 Pro, Gemini 2.5 Pro, and Gemini 2.5 Flash. RT was measured across nine representative biological tasks spanning three complexity tiers (simple, complex, and interactive), reflecting increasing levels of reasoning and execution requirements. For each model–task combination, 20 independent trials (n = 20) were conducted to ensure statistical robustness.

RT values are reported as mean ± s.e.m., with individual replicates visualized as jittered points to capture variability. A five-level sequential color gradient was used to represent mean RT across tasks, facilitating direct comparison of latency profiles across different model configurations.

## Supporting information

Associated files for SI

## Data availability

Native RNA nanopore sequencing datasets for HEK293T and HeLa cell lines datasets are publicly available and can be accessed from the ENA database under accession number PRJEB80229.

ONT benchmarking oligo datasets and corresponding sequence reference were downloaded from: https://epi2me.nanoporetech.com/rna-mod-validation-data.

The two datasets, namely WT HeLa and HeLa *TRUB1* KO, have been deposited in the NCBI Sequence Read Archive under accession PRJNA1459519.

## Code availability

Our source code is available at https://github.com/wanunulab/NanoCortex.

## ACKNOWLEDGEMENTS

We thank the Northeastern High Performance Computing team for their support. We acknowledge support by the National Institutes of Health (R01HG013304, M.W., and 5R01HG012856, S.R.).

## AUTHOR CONTRIBUTIONS

Q.X., Z.W., M.W. and S.H.R. conceived the idea. Q.X. led the development of the agent-based framework, with contributions from Z.W. M.S. provided the HeLa KO dataset for method evaluation, and benchmarked the NanoCortex platform. M.W. and S.H.R. supervised the project.

## COMPETING INTERESTS

The authors declare no competing interests.

## Notes

https://www.ncbi.nlm.nih.gov/bioproject/PRJNA1459519/

### Competing Interest Statement

The authors have declared no competing interest.

