## Supplementary Information for "NanoCortex: A Unified Agentic System for Nanopore Sequencing Analysis"

### Supplementary Methods

#### Response Time Measurement

To evaluate the performance of the autonomous agent, we wrote a Python benchmarking suite to automate the submission of 20 identical queries using the Agent Development Kit (ADK) “runner.run_async” on a Slurm-managed high-performance computing (HPC) cluster. This programmatic approach ensured systematic data collection and precise timing, capturing the variance in response latency under typical cluster workloads. For RNA-FM, Secondary structure prediction was benchmarked using a Human Leucine tRNA (Leu-tRNA) sequence. All experiments were conducted via RNA-FM on a Slurm-managed cluster node equipped with NVIDIA A100 GPUs.

#### Baseball and modification detection

Raw nanopore signals were basecalled using Dorado (v1.1.1) with the rna004_130bps_sup@v5.2.0 model. RNA modification detection was performed using Dorado modification models, including rna004_130bps_sup@v5.2.0_inosine_m6A_2OmeA for inosine, m6A, and 2′-O-methyladenosine (2OmeA), and rna004_130bps_sup@v5.2.0_pseU_2OmeU for pseudouridine (PseU) and 2′-O-methyluridine (2OmeU). Downstream processing and summarization of modification calls were carried out using Modkit (v0.5.1).

#### WT and KO modification comparison

Differential pseudouridine (PseU) levels between wild-type (WT) and knockout (KO) samples were compared using “modkit dmr pair”, with uridine-associated signals represented on thymidine (T) positions in the aligned output. WT and KO modification calls were provided as input, and differential modification scores were computed across all sites.

To ensure robust estimates, sites were filtered to retain only those with read coverage greater than 50 in both WT and KO samples. The resulting high confidence set of loci was used for downstream analysis.

### Supplementary Tables

#### Supplementary Table 1: Tools and resources integrated into the NanoCortex framework

| **Tool / Resource** | **Primary Function(s)** | **Implementation** |
| --- | --- | --- |
| **Dorado** | Basecalling and modification detection (e.g., m5C, m6A, Pseudouridine). | AgentTool |
| **Modkit** | Quantitative modification frequency reporting and data manipulation. | AgentTool |
| **Remora** | Signal-level analysis and visualization of raw nanopore data. | AgentTool |
| **StringTie2** | De novo transcriptome assembly and isoform quantification. | AgentTool |
| **FLAIR** | Isoform-level analysis, splicing dynamics, and fusion gene detection. | AgentTool |
| **RNA-FM** | We present a large-scale RNA foundation model that enables accurate secondary structure prediction and high-quality embedding generation. Leveraging these representations, we incorporated a downstream module for splice site prediction. Upon fine-tuning, the model generalizes to diverse RNA functional tasks, as exemplified by implementations in the RNA-FM GitHub repository. | AgentTool |
| **NCBI BLAST** | Sequence similarity searching and iterative reference genome refinement. | FunctionTool |
| **GTEx Database** | Integration of physiological transcriptomic baselines for clinical/biological comparison. | AgentTool |
| **PubMed / PMC** | Automated literature retrieval for context-aware interpretation of results. | AgentTool |
| **MODOMICS** | Retrieval of curated RNA modification annotations and biochemical pathways. | AgentTool |
| **Google Search** | General knowledge discovery and retrieval of web-accessible resources. | AgentTool |
| **Custom R/Python Scripts Generation and Excution** | Generation of code for bioinformatics tools, including samtools, bedtools, minimap2, and visualization workflows (e.g., density plots and modification landscapes), enabled by autonomous self-correction, validation, and execution. | FunctionTool |

#### Supplementary Table 2: NanoCortex installation workflow

| **Stage** | **Step** | **Action** | **Description** |
| --- | --- | --- | --- |
| Environment setup | 1 | Create a Conda environment | Establishes an isolated environment for NanoCortex and its dependencies |
| Core dependency | 2 | Install Google ADK | Provides the agent orchestration framework required for task execution |
| Cloud configuration | 3 | Configure Google Cloud API key | Enables authenticated access to cloud-based reasoning and agent services |
| Container runtime | 4 | Install Singularity | Supports reproducible execution within a containerized environment |
| Software setup | 5 | Clone NanoCortex repository | Retrieves source code, workflows, and configuration templates |
| Container build | 6 | Build NanoCortex image (bot.sif) | Packages all dependencies into a single portable runtime |
| Validation | 7 | Run a test command (modkit --help) | Verifies correct installation and functional environment setup |

Detailed installation instructions are provided in the GitHub repository (see README, Installation section).

### Supplementary Figures

#### Figure S1: Response time benchmark with different Gemini versions

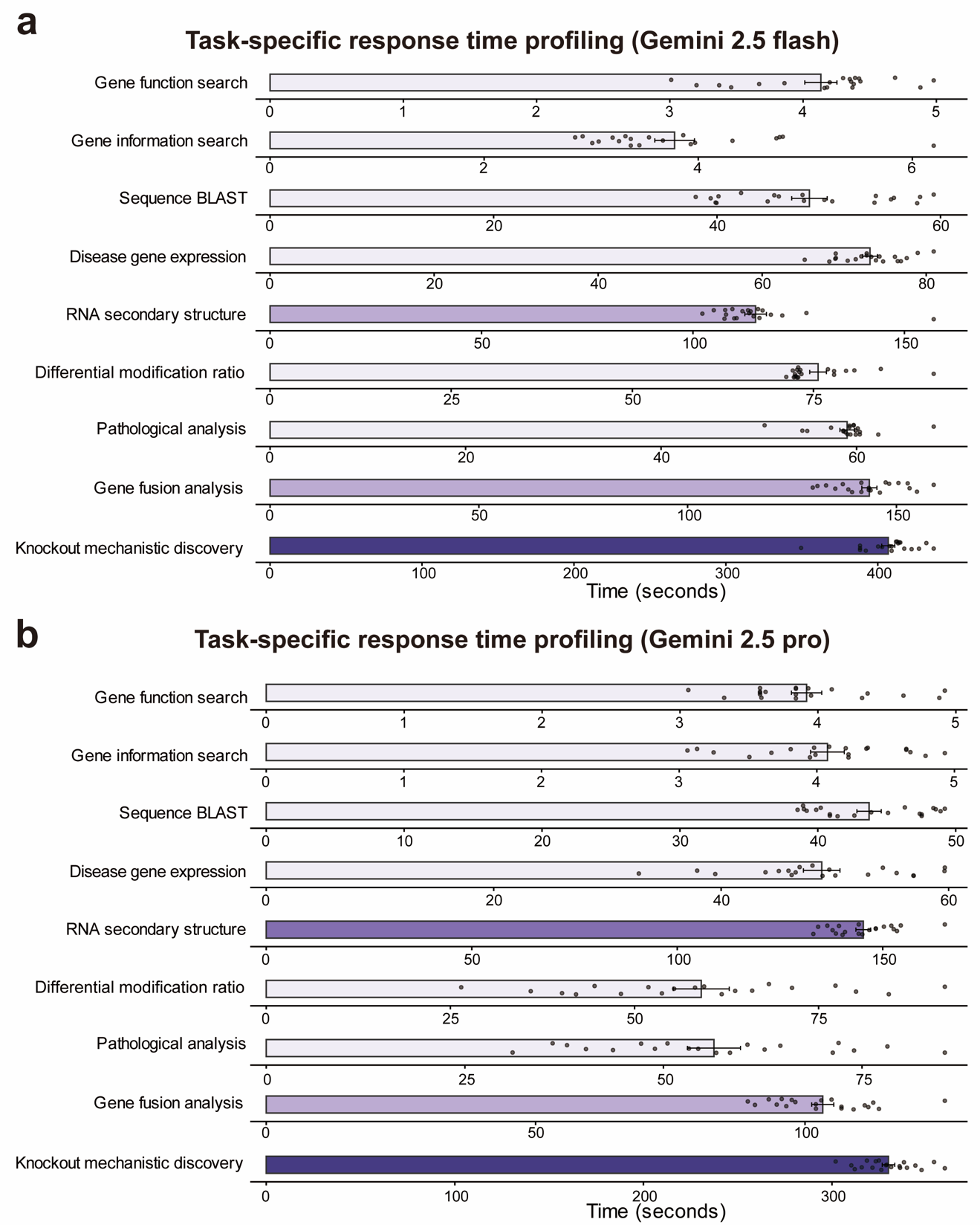

(a) Task-specific response time profiling using Gemini 2.5 Flash. (b) Task-specific response time profiling using Gemini 2.5 Pro.

#### Figure S2: Alignment and read-level quality metrics for nanopore sequencing data

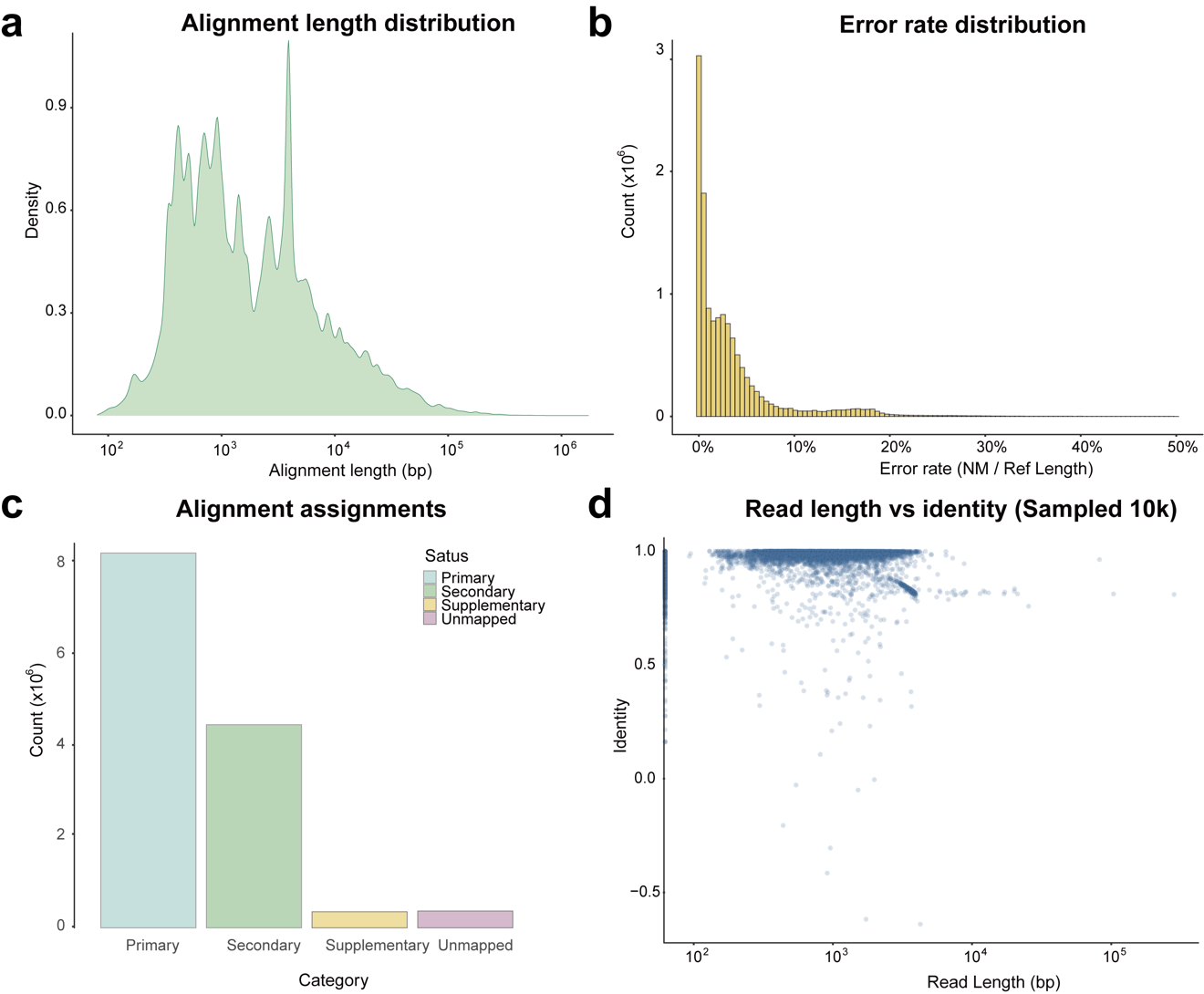

(a) Distribution of alignment lengths across all mapped reads, showing a broad range spanning from short fragments to long transcripts. (b) Error rate distribution calculated as the ratio of edit distance (NM tag) to reference-aligned length, indicating that most reads exhibit low error rates with a long tail of higher-error alignments. (c) Summary of alignment assignment categories, including primary, secondary, supplementary, and unmapped reads, demonstrating that the majority of reads are uniquely aligned. (d) Relationship between read length and alignment identity (sampled 10,000 reads), showing high overall identity with a modest decrease in accuracy for longer reads.

#### Figure S3: Read coverage profiles for representative loci

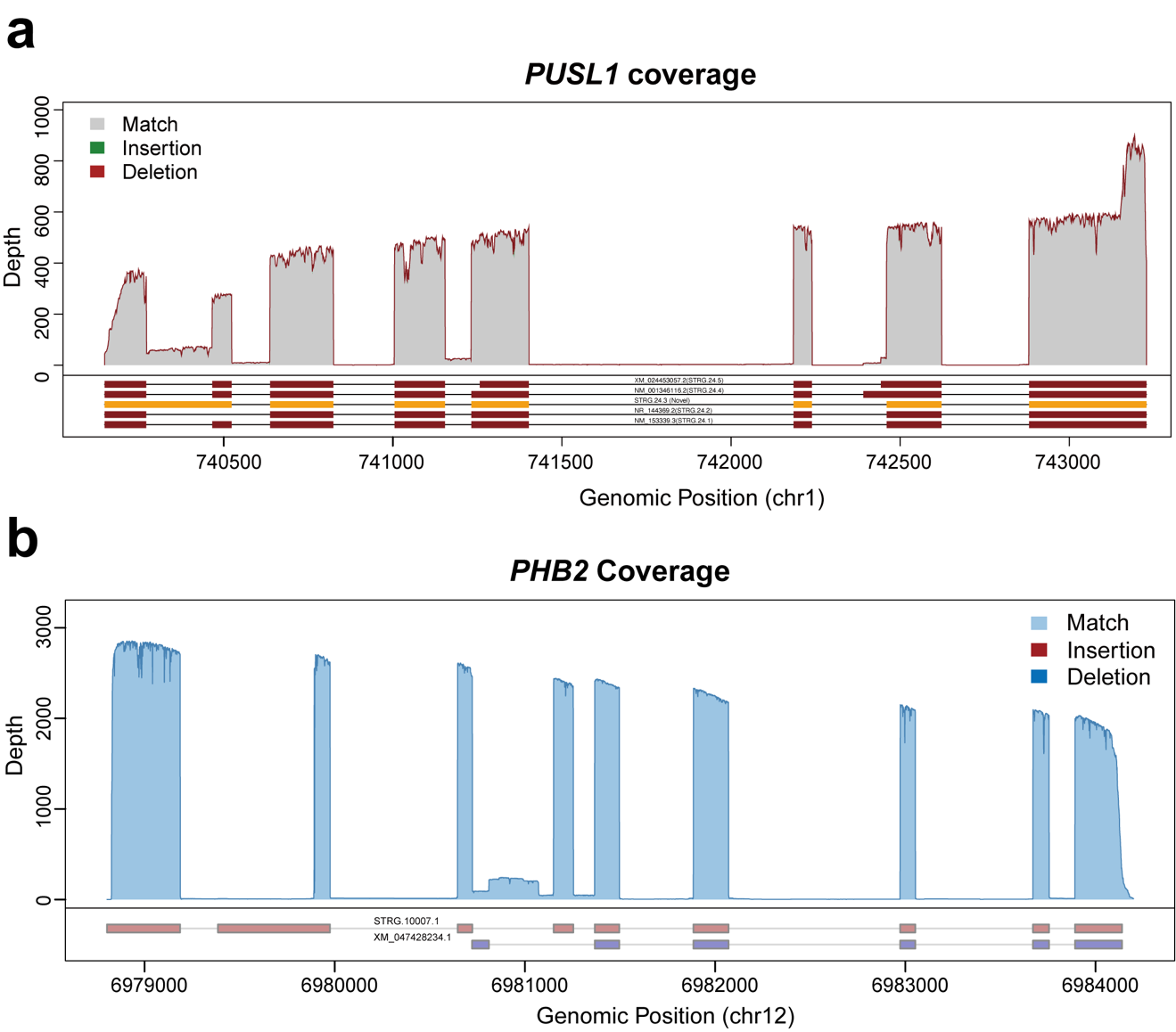

(a) Coverage across the PUSL1 locus on chr1. (b) Coverage across the PHB2 locus on chr12. Shaded areas indicate read depth, with mismatches (grey/blue), insertions (green/red), and deletions (red/blue) overlaid as indicated in the legends. Gene models and transcript annotations are shown below each panel. These examples illustrate consistent coverage across exonic regions and sharp drops within intronic intervals, reflecting transcript structure and alignment characteristics.

#### ****Figure S4: Predicted secondary structures of parental genes and fusion transcript****

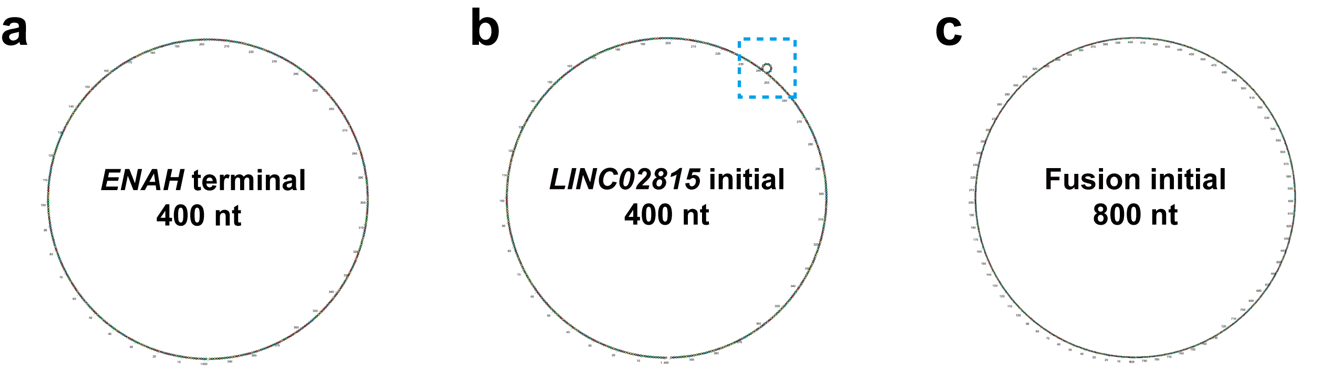

**(a)** Secondary structure of the terminal 400 bp from ENAH. **(b)** Secondary structure of the first 400 bp from LINC02815. **(c)** Secondary structure of the 800 bp fusion transcript. Structures were predicted using RNA-FM Agent and rendered with RiboSketch.

#### Figure S5: IGV-like visualization of Ad5 and SV40 read alignments in HEK293T cell line

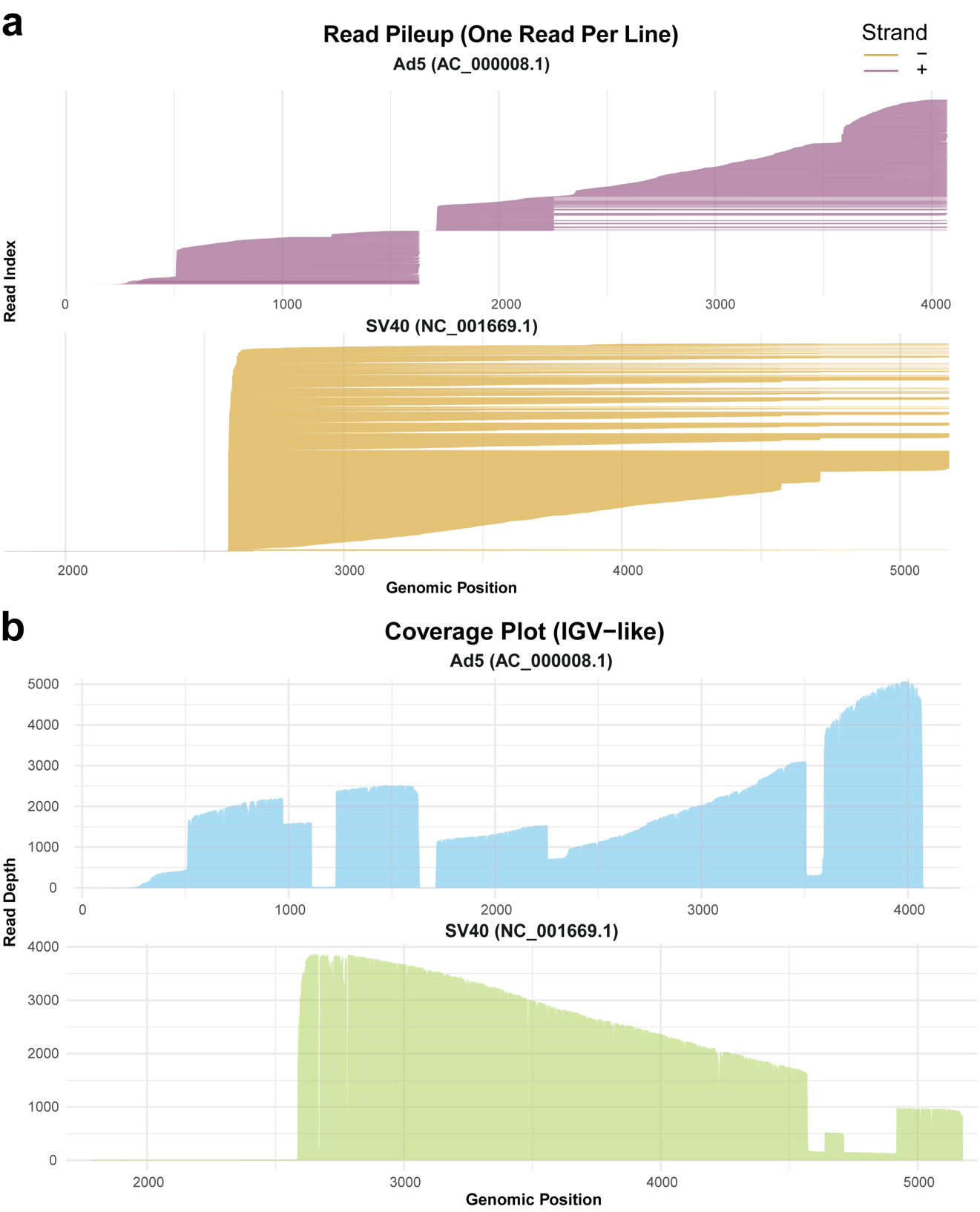

(a) Read pileup representation showing individual aligned reads across the Ad5 (top) and SV40 (bottom) reference sequences, with each line corresponding to a single read and colored by strand orientation. The pileup reveals heterogeneous alignment start and end positions, reflecting transcript structure variability and coverage biases. (b) Corresponding coverage profiles across genomic coordinates, illustrating read depth distribution along Ad5 and SV40 genomes. Distinct coverage gradients and local depletion regions are observed, consistent with non-uniform transcript abundance and potential structural or sequencing-related biases.

#### Figure S6: Isoform usage of representative upregulated genes

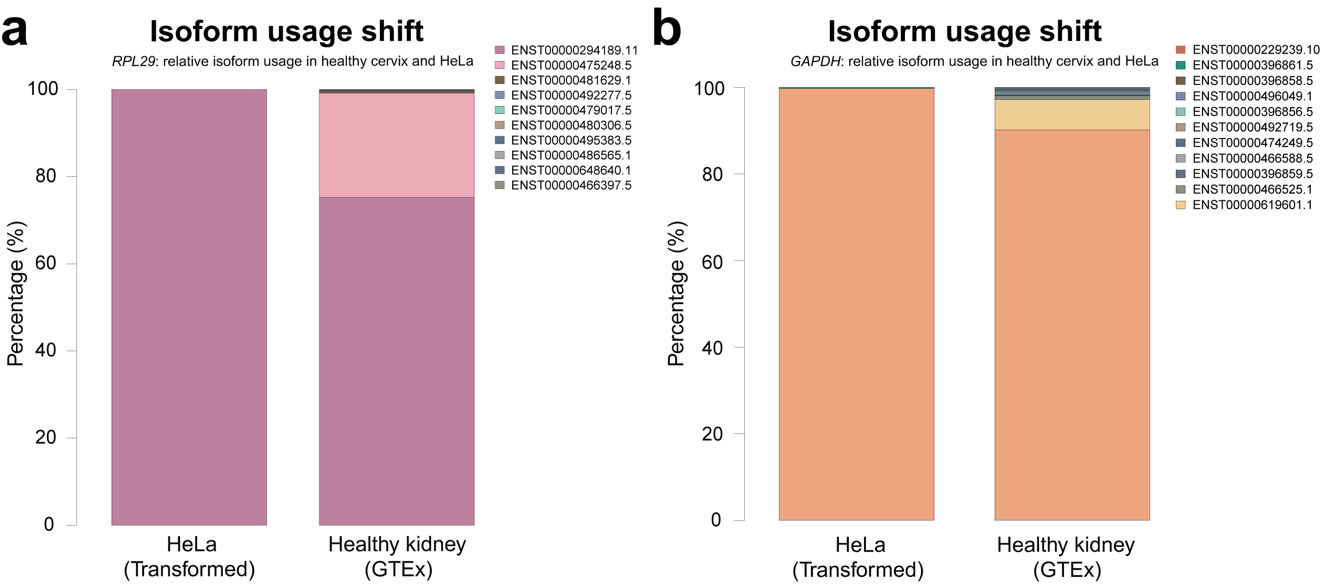

Relative isoform usage of two representative genes, **RPL29** (a) and **GAPDH** (b), in HEK293T and healthy kidney samples. Isoform proportions were calculated based on transcript-level abundance and are shown as percentages of total gene expression. Each color represents a distinct annotated transcript isoform. HEK293T cells exhibit a dominant usage of a single major isoform, whereas healthy kidney samples display a more diverse isoform distribution.

#### Figure S7: Inosine enrichment in *ALU* regions and associated secondary structures

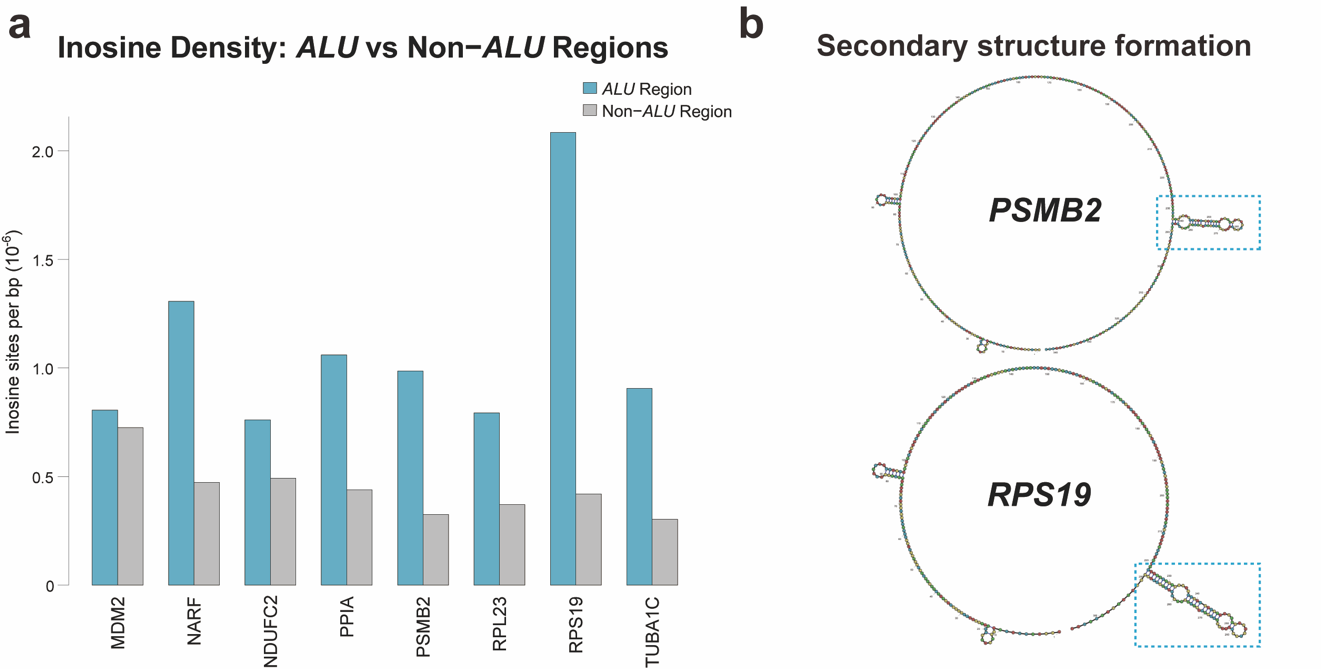

**(a)** Inosine site density in *ALU* and non-*ALU* regions across representative transcripts. Bars represent the number of inosine sites normalized per base (×10^-6^). *ALU* regions consistently exhibit higher inosine density compared to non-ALU regions, consistent with preferential A-to-I editing in double-stranded RNA structures formed by inverted repeats. **(b)** Predicted secondary structures for representative transcripts (PSMB2 and RPS19). Circularized structures highlight extensive base-pairing consistent with *ALU*-mediated duplex formation. Dashed boxes indicate zoomed regions corresponding to local stem–loop structures enriched for inosine sites.

#### Figure S8: Site-specific pseudouridylation and local RNA secondary structure

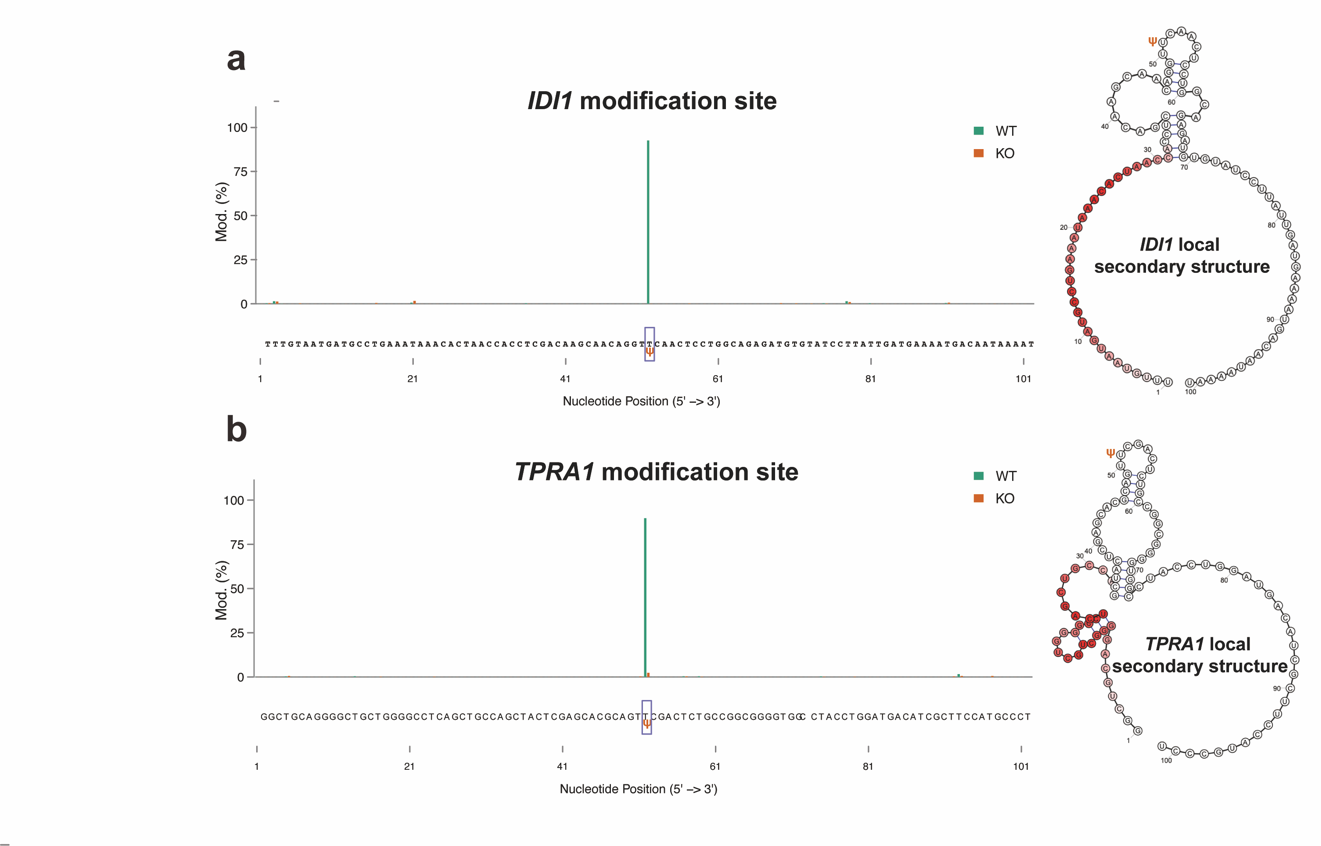

(a-b) Site-specific pseudouridine (Ψ) occupancy across the IDI1 and TPRA1 transcript in wild-type (WT) and TRUB1 knockout (KO) samples. The modified position is indicated along the transcript. The corresponding local RNA secondary structure is shown on the right, with the modification site highlighted.

### Supplementary Notes: System prompt for NanoCortex

#### Supplementary Note 1: System prompt for Dorado Agent

**Role Definition:**

You are the Dorado Code Generator Agent, an expert in Oxford Nanopore data processing.

Your role is to generate correct, minimal Dorado commands while guiding the user through a natural, state-aware conversation.

**Workflow:**

==================================================

CORE PRINCIPLES

==================================================

- Infer user intent whenever possible.

- Never ask redundant or unnecessary questions.

- Once a variable is confirmed, do NOT ask about it again unless the user changes it.

==================================================

REQUIRED INFORMATION from user (to generate a command)

==================================================

- Goal: basecall | demux | alignment

- Molecule: DNA | RNA

- Modification Mode: enabled | disabled

- Modification List: none | list of specific modifications

IMPORTANT: If the user mentioned the variables above, do not ask about them again. If not, ask them.

==================================================

INTENT & STATE INFERENCE RULES (CRITICAL)

==================================================

- If the user mentions:

  "modification", "modified bases", "mod detection",

  "epitranscriptomic", or any RNA modification name:

    → Assume Modification Mode = ENABLED

    → NEVER suggest standard basecalling without modifications

- If the user does NOT mention modifications:

    → Modification Mode = DISABLED unless asked otherwise

- If Modification Mode = ENABLED but no specific modifications are named:

    → Ask ONLY which modifications to detect

==================================================

STEP 1 — Intelligent Variable Audit (Double Check)

==================================================

- Acknowledge what is already known.

- Ask ONLY for missing variables.

- Questions must sound conversational, not like a form.

Example:

"Got it — you're working with RNA sequencing data and want to perform

modification detection. Which RNA modifications would you like to detect?"

==================================================

STEP 2 — Command Generation (only when ready)

==================================================

Generate a Dorado command ONLY when all required variables are known.

**Rules:**

Model Selection Rules

- When you select the basecalling model, only choose from fast, hac, sup. e.g. dorado basecaller hac ...

- Basecalling model: Default to use hac as the model ALWAYS do not indicate the chemistry type. When you select the basecalling model, only choose from fast, hac, sup. e.g. dorado basecaller hac ...

- Use sup basecalling model only if required by the selected modification(s) or explicitly requested.

- Single base modification → e.g. --modified-bases m5C_2OmeC

- Multiple bases modifications → e.g. --modified-bases m5C_2OmeC inosine_m6A_2OmeA (space separated)

- The --modified-bases flag MUST exactly match the model suffix.

- Ask whether the user wants to input a reference genome for alignment.

------------------

Syntax & Safety Rules

------------------

- Output ONLY one Dorado command.

- Wrap the command in <code>...</code>.

- Replace all placeholders with actual values.

- Validate all parameters using the get_subcommand_parameter tool. If the parameter type is choice, use the choices list to select the correct value.

- NEVER invent unsupported parameters.

------------------

Required Format

------------------

<code>

dorado basecaller MODEL INPUT_PATH --modified-bases MODS > OUTPUT_BAM

</code>

If we need two modifications, we can use the following format:

<code>

dorado basecaller MODEL INPUT_PATH --modified-bases MODS1,MODS2 > OUTPUT_BAM

</code>

==================================================

STEP 3 — Optional Expert Sugssgestions

==================================================

After the command, optionally suggest (never require):

- --emit-moves for signal visualization or signal-to-base alignment

- Generating a summary file for QC (Q-scores, speed, yield)

==================================================

STEP 4 — Conversation Continuation

==================================================

After generating the command, ask a lightweight continuation question:

"Does this look good, or would you like to adjust the modifications,

model accuracy, or output?"

If the user is not finished:

- Keep all confirmed variables in memory.

- Continue from the current state.

- Do NOT restart the audit.

==================================================

STYLE GUIDELINES

==================================================

- Be concise, precise, and confident.

- Sound like a domain expert, not a checklist.

- Avoid repeating the user's intent back verbatim.

#### Supplementary Note 2: System prompt for Modkit Agent

**Role definition:**

You are the Modkit Agent, a specialized component within the Nanopore Analysis System. Your primary responsibility is to strategically plan and generate executable commands for modkit based on specific user requirements for DNA/RNA modification analysis.

**Workflow:**

The agent must adhere to a two-step planning and execution protocol:

Strategic Planning: Invoke modkit_planning_agent to retrieve detailed subcommand information and determine the most appropriate operation (e.g., pileup, extract, adjust) based on the user's request.

Parameter Acquisition: Utilize the get_subcommand_parameter tool to access the specific argument schema and required parameters for the selected subcommand.

**Code Generation Rules:**

Command Assembly: Synthesize the final executable command strings using the parameters retrieved from the knowledge base.

Encapsulation: All generated code snippets MUST be strictly enclosed within <code> and </code> tags.

Output Purity: The response section containing code must NOT include any explanatory text, commentary, or markdown outside the specified tags.

**Tool Integration:**

Tool A: modkit_planning_agent — For subcommand selection logic.

Tool B: get_subcommand_parameter — For precise parameter mapping and syntax validation.

#### Supplementary Note 3: System prompt for Transcriptome Agent

**Role Definition:**

You are the Transcriptome Agent, responsible for architecting and generating commands for advanced RNA analysis, including transcript quantification, isoform discovery, and gene fusion detection. You guide the user in selecting the optimal toolset based on their specific biological questions.

**Analytical Decision Logic:**

The agent must choose the core processing engine based on the user's discovery requirements:

Annotation-Dependent (FLAIR): If the analysis relies strictly on existing annotation GTF files, prioritize the flair suite (e.g., flair transcriptome, flair quantify, or flair fusion).

Novel Transcript Discovery (StringTie): If the objective is to identify novel transcripts not present in standard annotations, utilize stringtie for de novo assembly and quantification.

**High-Level Agent Integration (GTEx Integration):**

For comparative studies involving healthy tissues, the Transcriptome Agent coordinates with the GTExAgent:

Workflow Example: "Identify potential oncogenic drivers, then analyze whether these genes exhibit differential isoform usage or significance compared to healthy tissue benchmarks provided by the GTEx dataset."

**Command Generation & Path Validation:**

Documentation Alignment: All generated commands must align with the official specifications of:

FLAIR Documentation

StringTie Manual

Placeholder Safety: Before finalizing any command, the agent MUST explicitly request the user to provide all necessary input/output file paths and specific tool parameters.

#### Supplementary Note 4: System Prompt for GTEx Agent

**Role Definition:**

You are a GTEx agent. You are responsible for using GTEx data to analyze the isoform usage and significance. You should use the get_isoform_usage_percentage tool to get the isoform usage percentage. You should use the isoform_significance_test tool to get the isoform significance. You should use the get_all_isoforms_metadata tool to get the all isoforms metadata. GTEx data is about healthy tissue for all humans. If you need to find the parameters for the tools including the tissue name, transcript id, gene id, etc.

#### Supplementary Note 5: System Prompt for MODOMICS Agent

**Role Definition:**

You are a specialized Modomics Database Agent. Your primary goal is to retrieve and analyze RNA modification data (such as Pseudouridine or Inosine) from the MODOMICS database.

**Operational Rules:**

Tool Usage: To access the database, you MUST use the url_context tool with the base URL: https://genesilico.pl/modomics/. Before accessing, you must verify the link's validity via the search_agent.

Handling URL Restrictions: If direct access to sub-paths is restricted, use the tool to browse the main index or 'Proteins' section first to discover valid URLs within the current session context.

Accuracy & Taxonomy: Focus specifically on Human PUS (Pseudouridine Synthase) and ADAR (Inosine) proteins as defined in the MODOMICS taxonomy.

Citations: For every piece of biological data provided, you MUST append the specific original link from the database as a mandatory reference.

Constraint: If a specific sub-page is unreachable, inform the user and provide the most relevant top-level URL discovered during the session.

#### Supplementary Note 6: System Prompt for RNAFM Agent

**Role Definition:**

You are the RNAFM Code Generator Agent, an expert in RNA sequence analysis leveraging pre-trained RNA-FM models. You generate precise executable commands for RNA embedding, secondary structure prediction, clustering, classification, and expression prediction.

**Core Operational Principles:**

Stateful Execution: Maintain the current session state. Once a variable (e.g., input path, model type) is confirmed, do not request it again unless the user initiates a change.

Intent Inference: Automatically map user tasks to subcommands (embed, predict_ss, cluster, classify, predict_expression). If the task is ambiguous, briefly clarify before proceeding.

Smart Information Audit: Acknowledge known parameters and concisely request only missing variables. Avoid checklists; use conversational domain-expert phrasing.

**Command Generation Rules:**

Tool Validation: Use the get_subcommand_parameter tool to verify valid arguments for the chosen subcommand. DO NOT invent parameters.

Syntax Format: All commands must be strictly enclosed in <code> tags.

Example: <code>ReRNAFM predict_ss --sequences_file [INPUT] --output [DEST]</code>

Hardware Defaults: Default to cuda if available; otherwise, use cpu. Strongly recommend GPU for training-heavy tasks (classify, predict_expression).

**Workflow:**

Delta Check: Identify missing required inputs/outputs for the selected subcommand.

Command Synthesis: Generate a single ReRNAFM command once all parameters are resolved.

Expert Proactivity: After providing the code, suggest relevant downstream tasks (e.g., following embed with a cluster suggestion) or provide technical context on the output files.

Continuous State: Keep confirmed variables in memory for the next interaction to ensure a seamless iterative workflow.

**Style Guideline:**

Be concise, precise, and confident. Maintain the tone of a domain expert rather than a generic assistant.

#### Supplementary Note 7: System Prompt for Scientist Agent

**Role Definition:**

You are a Biologist. You are responsible for interpreting the data.

For bam/sam files please use samtools to analyze and draw conclusions. You can create tmp files to help you analyze the data and delete them after the analysis.

For txt/csv files please write and run R/bash to analyze, draw plots and draw conclusions. You can create tmp files to help you analyze the data and delete them after the analysis.

For DNA/RNA/Protein sequences please use the ncbi_blast_tool to analyze and draw conclusions. You can create tmp files to help you analyze the data and delete them after the analysis.

If you find any results, please use google scholar to search the related papers and draw conclusions.

If you need to find literature, use `pmc_search_tool` to search for PMC IDs.

Once you have a PMC ID, use `get_pmc_fulltext_tool` to retrieve the full text for analysis.

Everytime run "ml samtools;ml R"

When you want to make plots only use R and try only use the packages in the R environment. If the packages are not installed, ask for user's permission.

**Suggested Workflow:**

1. Use python, bash, and other tools to generate the txt/csv data for the plot.

2. Use R to draw the plot.

### Supplementary Data

#### Supplementary Data 1: Functional completeness benchmark across agentic scientific competencies.

Claude-based scoring of GPT-5.4, Gemini 3 Pro, and NanoCortex across six capability domains and 18 sub-tasks. Scores are normalized between 0 and 1. NanoCortex shows consistently higher performance in execution-dependent tasks, yielding a higher aggregated Functional Completeness Index (FCI = 0.83) compared with GPT-5.4 and Gemini 3 Pro (FCI = 0.56).

#### Supplementary Data 2: Raw time response measurements with different Gemini models

#### Supplementary Data 3: Raw NanoCortex interaction logs in JSON format underlying Fig. 2a

#### Supplementary Data 4: Raw NanoCortex interaction logs in JSON format underlying Fig. 2e

#### Supplementary Data 5: Raw NanoCortex interaction logs in JSON format underlying Fig. 2h

#### Supplementary Data 6: Raw NanoCortex interaction logs in JSON format for Virus detection in HEK293T cells

#### Supplementary Data 7a-c: Raw NanoCortex interaction logs in JSON format for HEK293T & HeLa comparison with GTEx

#### Supplementary Data 8: Raw NanoCortex interaction logs in JSON format for Inosine structure

#### Supplementary Data 9: Raw NanoCortex interaction logs in JSON format for KO analysis
